# Characterization of three-dimensional bone-like tissue growth and organization under influence of curvature and directional fluid flow

**DOI:** 10.1101/2022.08.18.504382

**Authors:** Bregje W.M. de Wildt, Feihu Zhao, Iris Lauwers, Bert van Rietbergen, Keita Ito, Sandra Hofmann

## Abstract

The transition in the field of bone tissue engineering from bone regeneration to three-dimensional *in vitro* models has come with the challenge of recreating a dense and anisotropic bone-like extracellular matrix with cell culture. The creation of such an organized bone-like extracellular matrix has received little attention thus far. Although the mechanism by which bone extracellular matrix gains its structure is not fully understood, curvature (especially concavities), mechanical loading due to deformations or directional fluid flow, and osteocyte signaling have been identified as potential contributors. Here, guided by computational simulations, we evaluated three-dimensional cell and bone-like tissue growth and organization in a concave channel with and without directional fluid flow stimulation. Human bone-marrow derived mesenchymal stromal cells were seeded on donut-shaped silk fibroin scaffolds and stimulated to undergo osteogenic differentiation for 42 days statically or in a flow perfusion bioreactor. Constructs were investigated for cell distribution, and tissue growth and organization on day 14, 28, and 42. As a result, directional fluid flow was able to improve bone-like tissue growth but not organization. After 28 days of culture, when osteogenic differentiation was likely accomplished, cells tended to have a small preference for orientation in the tangential (*i.e*., circumferential) direction of the channel. Based on our results, we suggest that three-dimensional bone-like tissue anisotropy might be guided by curvature, while extracellular matrix production can be increased through the application of fluid shear stress. With this study, an initial attempt in three-dimensions was made to improve the resemblance of *in vitro* produced bone-like extracellular matrix to the physiological bone extracellular matrix.

## 1. Introduction

Bones have remarkable mechanical properties due to their extracellular matrix (ECM) composition and organization. To attain these properties, organic and inorganic matrix components are highly organized (Reznikov et al., 2014). In addition, bone structure is maintained and adapted through lifelong remodeling by osteoclasts (bone-resorbing cells), osteoblasts (bone-forming cells), and osteocytes (regulating cells) (Bonewald, 2011; Strandring et al., 2005). Traditionally, bone tissue engineering methods (*i.e*., making use of cells, scaffolds, biochemical and biomechanical stimuli) have been applied to recreate bone-like tissue *in vitro* for implantation and subsequent regeneration of large osseous defects. Because of bone’s innate remodeling capacity, these implants may successfully induce regeneration, even if they fail to mimic the complex bone ECM structure. The recapitulation of the physiological bone ECM *in vitro* has received too little attention from researchers (de Wildt et al., 2019).

Tissue engineering strategies are nowadays increasingly applied for the creation of *in vitro* models of healthy or pathological bone, aiming at improving preclinical treatment development while addressing the principle of reduction, refinement, and replacement of animal experiments (3Rs) (Balls et al., 1995; Caddeo et al., 2017; Holmes et al., 2009; Owen and Reilly, 2018). Changes in bone’s ECM composition and organization are characteristic for bone pathologies like osteoporosis, osteogenesis imperfecta, and bone metastasis (Bala and Seeman, 2015; Bishop, 2016; Karunaratne et al., 2016; Matsugaki et al., 2021). Therefore, *in vitro* models that aim at studying changes in bone ECM under the influence of treatments would benefit from improved control over organic matrix formation and subsequent mineralization (Hadida and Marchat, 2020; de Wildt et al., 2019). As the organic bone ECM with mainly collagen type 1 functions as a mineralization template (Wang et al., 2012), the improvement of collagen network organization and density may enhance the biomimicry of *in vitro* produced bone ECM. However, the mechanism by which collagen forms a dense anisotropic or lamellar network *in vivo* is poorly understood. It is well accepted that *in vivo* bone morphology and mass is regulated by osteocytes which sense interstitial fluid flow through their lacuna-canalicular networks (Burger and Klein-Nulend, 1999). Recently, the anisotropy of the osteocyte lacuna-canalicular network has been correlated with the degree of apatite orientation in bone ECM, indicating a role for osteocytes in regulating ECM anisotropy (Ishimoto et al., 2021). The preferred orientation of collagen producing osteoblasts could also be manipulated to stimulate anisotropic collagen formation. This might be accomplished by mechanical loading like cyclic stretch or directional fluid flow (Matsugaki et al., 2013; Xu et al., 2018). These studies are however mainly performed in a controlled but simplified two-dimensional (2D) environment for a short period of time, not representative for the *in vivo* situation and ignoring tissue formation outcomes. In three-dimensional (3D) systems, fluid flow has been demonstrated to stimulate bone-like tissue growth including collagen formation (Akiva et al., 2021; Melke et al., 2018; Vetsch et al., 2016). One challenge in these 3D environments is that both increased mass transport and wall shear stress (WSS) as a result of fluid flow could have an effect (Hadida and Marchat, 2020; Wittkowske et al., 2016).

In addition to the application of external mechanical loading, substrate curvature could also induce cell organization (Callens et al., 2020, BioRxiv), subsequent anisotropic collagen formation (Bidan et al., 2012), and bone-like tissue growth (Ehrig et al., 2019; Rumpler et al., 2008; Vetsch et al., 2016). Bone-like tissue growth is especially stimulated in concavities with high curvatures (Callens et al., 2020a). Cell and tissue anisotropy is then often observed in the tangential or circumferential direction of a pore (Bidan et al., 2012). Thus, to promote *in vitro* bone-like tissue growth and anisotropy, fluid flow and curvature are likely two important factors. However, to our knowledge these two factors have not been evaluated together and therefore it is unclear which of the two factors dominates bone-like tissue growth and anisotropy.

Accordingly, in this study bone-like tissue growth and anisotropy were evaluated in a concave channel in a 3D silk fibroin (SF) scaffold statically or under influence of directional fluid flow. Osteogenically stimulated human bone marrow-derived mesenchymal stromal cells (hBMSCs) were used because of their ability to proliferate and differentiate into osteoblasts and osteocytes (Akiva et al., 2021), making them an appropriate candidate for (personalized) *in vitro* bone models (Ansari et al., 2021). As the cell response to fluid flow or curvature might change during osteogenic differentiation from mesenchymal stromal cell to osteocyte (Callens et al., 2020a; Wittkowske et al., 2016), tissue growth and organization were studied over a period of 42 days with intermediate time points at day 14 and day 28 (Figure 1). In addition, prior to experiments computational simulations were performed. Fluid flow patterns and fluid shear stress magnitude at the channel wall were simulated with a computational fluid dynamics (CFD) model to i) determine the optimal bioreactor settings for osteogenesis and bone-like tissue formation, and ii) ensure only fluid flow at the channel wall in the longitudinal direction (*i.e*,. direction of the flow) to minimize an effect of mass transport and flow in the radial (*i.e*., into the scaffold) direction.

**Figure 1.**
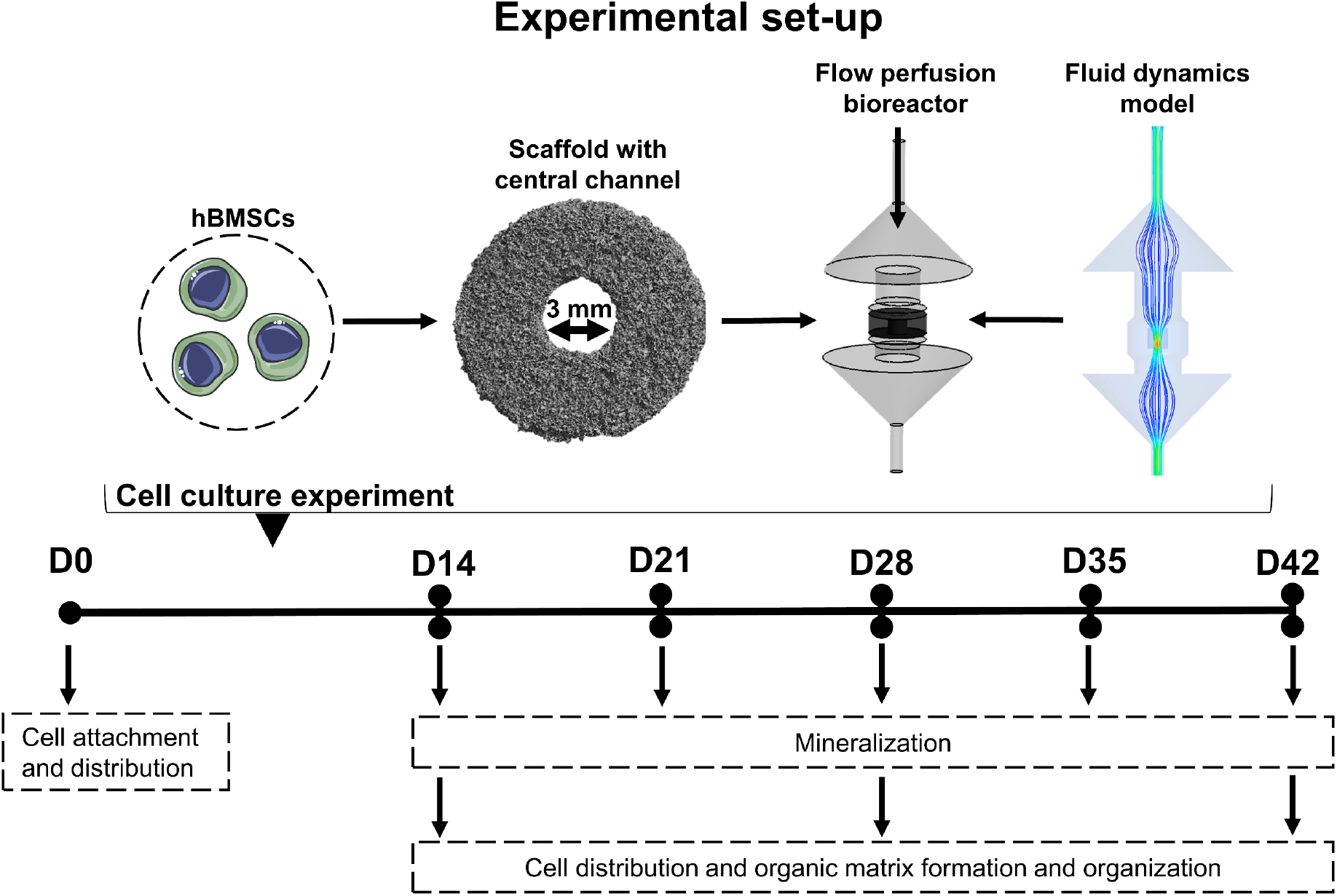
Experimental set-up of the study. hBMSCs were seeded on silk fibroin scaffolds with a central concave channel. Fluid flow was applied in longitudinal direction of the channel with a flow perfusion bioreactor of which the settings were determined with a computational fluid dynamics model. Cells were stimulated to undergo osteogenic differentiation over a period of 42 days with intermediate time points at day 14 and 28 to study cell distribution, and tissue growth and organization. Mineralization was checked weekly from day 14 onwards with non-destructive micro-computed tomography scanning. Abbreviations: human bone marrow derived stomal cells (hBMSCs), day (D). Cell images were modified from Servier Medical Art, licensed under a Creative Common Attribution 3.0 Generic License (http://smart.servier.com/, accessed on 8 July 2021).

## 2. Materials and Methods

### 2.1 Scaffold fabrication

Bombyx mori L. silkworm cocoons were degummed by boiling them in 0.2 M Na2CO3 (S-7795, Sigma-Aldrich, Zwijndrecht, The Netherlands) for 1 h. Silk was dried and subsequently dissolved in 9 M LiBr (199870025, Acros, Thermo Fisher Scientific, Breda, The Netherlands), filtered, and dialyzed against ultra-pure water (UPW) for 36 h using SnakeSkin Dialysis Tubing (molecular weight cut-off: 3.5 K, 11532541, Thermo Fisher Scientific). Dialyzed SF solution was frozen at −80° C and subsequently lyophilized for 6 days. Lyophilized SF was dissolved in hexafluoro-2-propanol (003409, Fluorochem, Hadfield, UK) at a concentration of 17% (w/v) and casted in cylindrical scaffold molds filled with NaCl granules with a size of <200 *μ*m as templates for the pores. Molds were covered and after 3 h, covers were removed from molds, and hexafluoro-2-propanol was allowed to evaporate for 7 days whereafter *β*-sheets were induced by submerging SF-salt blocks in 90% MeOH for 30 min. SF-salt blocks were cut into discs of 3 mm height with a Accutom-5 (04946133, Struer, Cleveland, OH, USA). NaCl was dissolved from the scaffolds in ultra-pure water, resulting in porous sponges. These sponges were punched with a 9 mm diameter biopsy punch for the outer dimensions and a 3 mm diameter biopsy punch for the central channel with a fixed curvature of −0.67 mm-1. The dimensions of the channel are based on previous research in which a 3 mm channel remained open over a period of 42 days (Vetsch et al., 2016), which is essential to enable studying the influence of directional fluid flow. Scaffolds were sterilized by autoclaving in phosphate buffered saline (PBS) at 121° C for 20 min.

### 2.2 Scaffold geometry

To obtain the detailed geometry of the scaffold, a micro-computed tomography scan (*μ*CT) was acquired with a *μ*CT100 imaging system (Scanco Medical, Brüttisellen, Switzerland). Scanning was performed for air-dried scaffolds with an isotropic voxel size of 3.9 *μ*m, energy level of 45 kVp, intensity of 200 *μ*A, integration time of 300 ms, and with twofold frame averaging. To reduce part of the noise, a constrained Gaussian filter was applied with a filter support of 1 and a filter width sigma of 0.8 voxel. Filtered images were segmented at a global threshold of 55% of the maximum grayscale value. Unconnected objects smaller than 50 voxels were removed through component labeling. A distance transformation function was used to determine the pore size distribution at four regions of interest (location S1, S3, S5, and S7 of Figure 2A).

**Figure 2.**
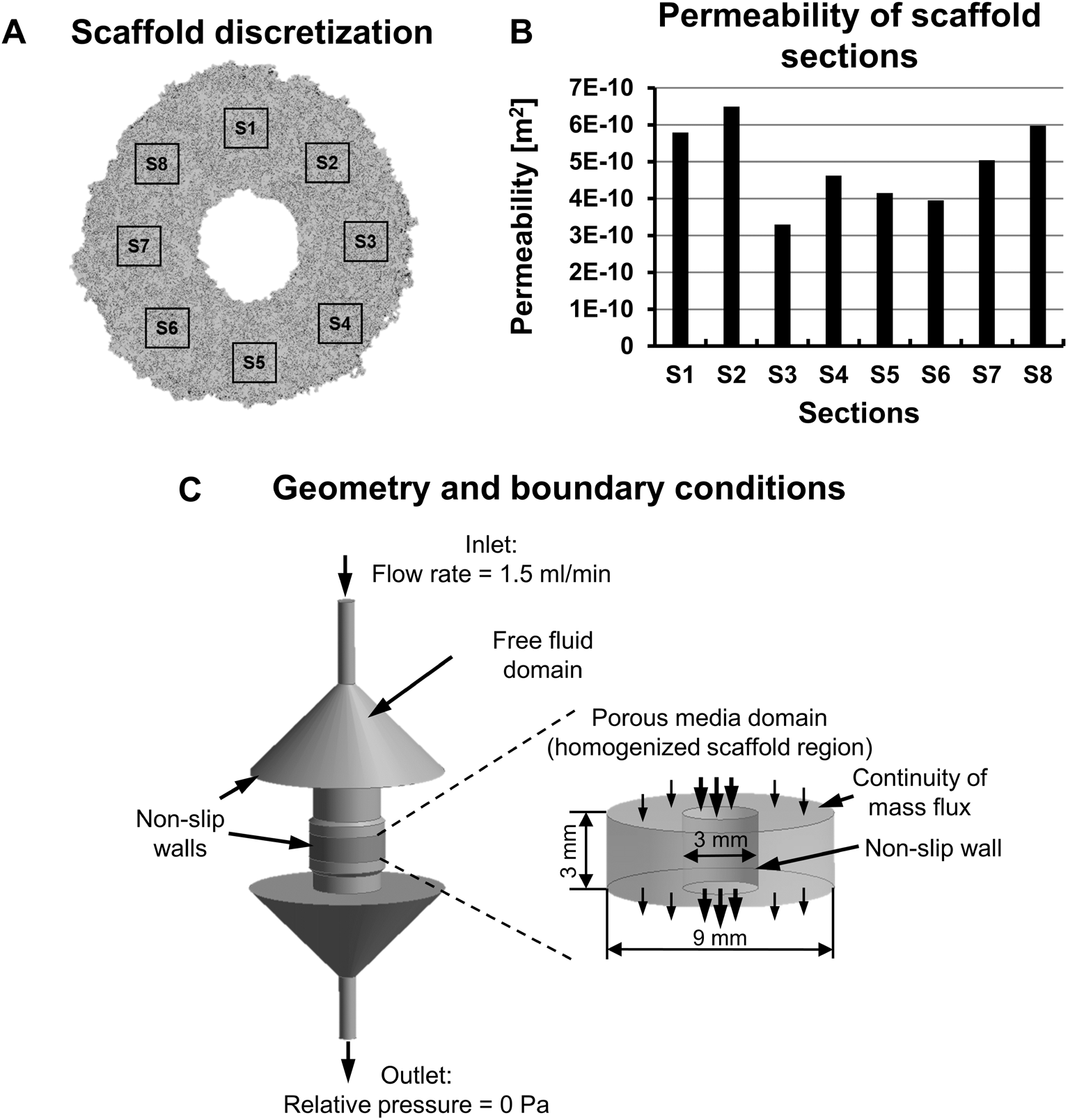
Multi-scale CFD model parameters. (**A**) Scaffold geometry was obtained with *μ*CT scanning whereafter the reconstructed scaffold was discretized into eight RVEs for permeability determination using the micro-model. (**B**) Calculated permeability of each scaffold RVE. (**C**) Geometry and boundary conditions of the CFD model. Abbreviations: computational fluid dynamics (CFD), representative volumetric element (RVE), micro-computed tomography (*μ*CT)

### 2.3 Multi-scale computational fluid dynamics (CFD) model

#### 2.3.1 Micro-scale: scaffold permeability calculation

As the region of interest was the scaffold’s central channel, the porous region around the channel was homogenized using the permeability that was determined based on the reconstructed geometry from the *μ*CT scan as previously described (Zhao et al., 2019). This was done to reduce the excessively high computational cost caused in modeling the irregular micro-struts of the porous SF scaffold. The homogenized scaffold permeability was determined based on the geometry of 8 representative volumetric elements (RVEs) with a diameter of 500 *μ*m (> 4 times average pore size) and a height equal to the height of the scaffold (Figure 2A). The RVEs were selected from the total scaffold geometry that was reconstructed using Seg3D software (University of Utah, UT, USA). The fluid domain of RVEs was meshed using the same strategy as in (Zhao et al., 2019) with global maximum and minimum element sizes if 20 *μ*m and 0.2 *μ*m, respectively.

The RVEs’ permeability was determined from Darcy’s law (Equation 1):

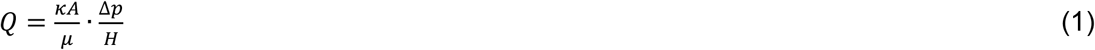

Where Δ*p* is the pressure drop over the scaffold height *H* determined by solving the CFD model for each RVE; *Q* is the prescribed flow rate, *A* the cross-sectional area to the flow, *μ* the dynamic viscosity of the culture medium (*μ* =1.09 mPa·s for cell culture medium (Maisonneuve et al., 2013)), and *κ* the permeability.

In the CFD model, the medium flow was defined as incompressible Newtonian and described by the Navier-Stokes equation that was solved using the finite volume method (FVM). As a convergence criterion it was required that the root-mean-square residual of the mass and momentum was smaller than a fixed threshold set at 10^-4^. Calculations were done using ANSYS CFX (ANSYS, Inc., PA, USA).

#### 2.3.2 Macro-scale: wall shear stress calculation

The macro-scale model representing the full scaffold and perfusion bioreactor domain was used for the fluid shear stress calculations on the scaffold channel wall (Figure 2C). In this macro-structural model, the scaffold region was homogenized and assigned a permeability calculated from the eight RVE micro-scale models. The region of the porous media (scaffold and newly formed tissue) was defined as a homogeneous porous media (Figure 2C), and described by Darcy’s equation:

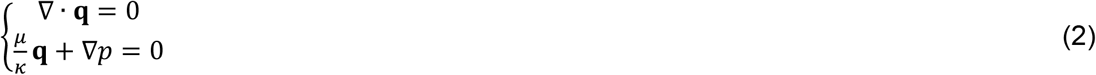

where **q** is the Darcy velocity and *p* the pressure.

The remaining central bioreactor channel without tissue formation was defined as free fluid (incompressible, Newtonian laminar flow), and described by the Navier–Stokes equations:

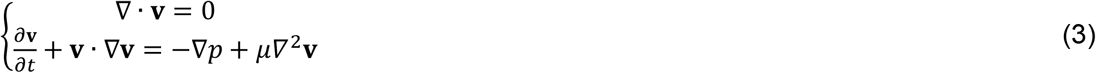

where **v** is the fluid velocity vector.

The top and bottom surfaces of the porous media domain were defined as boundaries with continuity of mass flux, and the scaffold internal channel wall was defined as a non-slip wall boundary as it is assumed that there will be only minor fluid following across the wall. At the inlet of the CFD model, a constant flow rate of 1.5 mL/min was prescribed according to the experimental condition. At the outlet, a relative pressure of 0 Pa was applied. The macro-scale CFD model was meshed with 1,720,090 tetrahedral elements, solved by FVM using ANSYS CFX (ANSYS, Inc., PA, USA) and the same convergence criteria as for micro-scale model.

To check the assumption that only little fluid flows through the interface between the channel and the scaffold, a CFD model in which a scaffold with idealized cubic pore shape and uniform pore size, was utilized. As this model only served to support the assumptions made for the main macro-model as described above, details on this model can be found in the supplementary information (Section 2).

### 2.4 Cell experiment

#### 2.4.1 hBMSC isolation, expansion and seeding

hBMSCs were isolated from human bone marrow (Lonza, Walkersville, MD, USA) and characterized for surface markers and multilineage differentiation, as previously described (Hofmann et al., 2007). hBMSCs were frozen at passage 4 with 5*10^6^ cells/ml in freezing medium containing fetal bovine serum (FBS, BCBV7611, Sigma-Aldrich) with 10% DMSO and stored in liquid nitrogen until further use. Before experiments, hBMSCs were thawed, collected in high glucose DMEM (hg-DMEM, 41966, Thermo Fisher Scientific), seeded at a density of 2.5*10^3^ cells/cm^2^ and expanded in expansion medium containing hg-DMEM, 10% FBS (BCBV7611, Sigma-Aldrich), 1% Antibiotic Antimycotic (anti-anti, 15240, Thermo Fisher Scientific), 1% Non-Essential Amino Acids (11140, Thermo Fisher Scientific), and 1 ng/mL basic fibroblastic growth factor (bFGF, 100-18B, PeproTech, London, UK) at 37 °C and 5% CO_2_. After 7 days, cells were detached using 0.25% trypsin-EDTA (25200, Thermo Fisher Scientific) and seeded onto scaffolds at passage 5. Cells were seeded at a density of 5*10^6^ cells per scaffold and seeding was performed dynamically for 6 hours in 50 ml tubes on an orbital shaker at 150 RPM in osteogenic control medium (lg-DMEM (22320, Thermo Fisher Scientific), 10% FBS (SFBS, Bovogen, East Keilor, Australia and 1% anti-anti) (32). After seeding, scaffolds were kept in wells plates overnight to ensure proper attachment before flow was applied.

#### 2.4.2 Bioreactor culture

hBMSC-loaded scaffolds were cultured in custom-made flow perfusion bioreactors as previously described (Vetsch et al., 2017). Scaffolds were cultured statically and dynamically for which a flow of 1.5 ml/min was applied, based on pre-simulations. Cells were stimulated to undergo osteogenic differentiation by providing osteogenic differentiation medium (osteogenic control medium with osteogenic supplements 10 mM β-glycerophosphate (G9422, Sigma-Aldrich), 50 μg/ml ascorbic acid-2-phosphate (A8960, Sigma Aldrich), and 100 nM Dexamethasone (D4902, Sigma-Aldrich)). For static bioreactors, 6 ml medium was supplied which was completely changed 3 times a week. For dynamic bioreactors, 12 ml medium was supplied of which only half of the volume could be replaced 3 times a week with double concentrated osteogenic supplements (*i.e*,. 20 mM β-glycerophosphate, 100 μg/ml ascorbic acid-2-phosphate, and 200 nM Dexamethasone) to keep the initial concentration of supplements constant. The bioreactor culture was maintained for 14, 28 or 42 days at 37 °C and 5% CO_2_.

### 2.5 Cell construct analyses

#### 2.5.1 Cell attachment

As cell attachment can be influenced by the cell experienced mechanical load (Mccoy and Brien, 2010), cell attachment was assessed at day 0 for a small subset of the samples (*N* = 4). These samples were cut in half and fixed in 2.5% glutaraldehyde in 0.1 M sodium cacodylate buffer (CB) for 4 h and then washed in CB. Samples were stained with 0.04% osmium tetroxide (75632, Sigma-Aldrich) in CB for 90 min and dehydrated with graded ethanol series (37%, 67%, 96%, 3 x 100%, 15 min each) followed by lyophilization. Samples were subsequently coated with 20 nm gold and imaging was performed in high vacuum, at 10 mm working distance, with a 10 kV electron beam (Quanta 600F, FEI, Eindhoven, The Netherlands).

#### 2.5.2 Cell distribution visualization

Day 0, 14, 28 and 42 scaffolds (*N* = 4 scaffolds per condition per time point) were cut in half and soaked for 15 minutes in each 5% (w/v) sucrose and 35% (w/v) sucrose in PBS. Samples were embedded in Tissue Tek^®^ (Sakura, Alphen aan den Rijn, The Netherlands) and quickly frozen with liquid N2. Cryosections were sliced in the vertical plane with a thickness of 5 *μ*m. Upon staining, sections were fixed for 10 minutes in 3.7% neutral buffered formaldehyde and washed twice with PBS. Sections were subsequently stained with 1 *μ*g/ml DAPI (D9542, Sigma-Aldrich) for 10 min and washed twice with PBS. Coverslips were mounted with Mowiol, tile scans were made with an epi-fluorescence microscope (Zeiss Axio Observer 7, 10x/0.3 EC Plan-Neofluor), and tile scans were stitched with Zen Blue software (version 3.3, Zeiss, Breda, The Netherlands).

#### 2.5.3 Organic matrix growth visualization and quantification

To visualize collagen deposition, cryosections were prepared in two different planes (*i.e*., the horizontal and vertical place, *N* = 4 scaffolds per condition, time point, and plane) and stained with Picrosirius Red. Sections were soaked in Weigert’s Iron Hematoxylin (HT1079, Sigma-Aldrich) solution for 10 minutes, washed in running tap water for 10 minutes, and stained in 1% w/v Sirius Red (36,554-8, Sigma-Aldrich) in picric acid solution (36011, Sigma-Aldrich) for one hour. Subsequently, sections were washed in two changes of 0.5% acetic acid and dehydrated in one change of 70% and 96% EtOH, three changes of 100% EtOH, and two changes of xylene. Sections were mounted with Entellan (107961 Sigma-Aldrich). To capture the entire section, tile scans were made with a bright field microscope (Zeiss Axio Observer Z1, 10x/0.45 Plan-Apochromat objective). Tile scans were stitched with Zen Blue software (version 3.1, Zeiss). Tissue growth was quantified in the vertical plane and horizontal plane measuring the distance from outer scaffold boarder to outer cell/tissue boarder at 10 different locations in each plane using Fiji (Schindelin et al., 2012). The measured distances in the two planes were averaged for the average channel tissue ingrowth per sample.

To check whether the produced ECM was of a bone-like nature, cryosections (*N* = 4 scaffolds per condition per time point) were prepared and fixed as described above (Section 2.4.2) and stained with the bone ECM markers osteopontin and collagen type 1 (Franz-odendaal et al., 2006; Rutkovskiy et al., 2016). Briefly, sections were permeabilized in 0.5% Triton X-100 in PBS for 5 min and blocked in 10% normal goat serum in PBS for 30 min. Primary antibodies (osteopontin: 1:200, 14-9096-82, Thermo Fisher Scientific; collagen type 1: 1:200, ab34710, Abcam, Cambridge, UK) were incubated overnight at 4 °C, secondary antibodies (osteopontin: Alexa 546, 1:200, A21123, Invitrogen Molecular Probes; collagen type 1: Alexa 647, 1:200, A21246, Invitrogen Molecular Probes) were incubated with 0.1 *μ*g/ml DAPI for 1 h at room temperature. Coverslips were mounted with Mowiol and images were acquired with a laser scanning microscope (Leica TCS SP8X, 40x/0.95 HC PL Apo objective).

#### 2.5.4 Biochemical content analyses

To quantify the biochemical content, lyophilized scaffolds (*N* = 4 per condition per time point) were weighted and digested overnight in papain digestion buffer (containing 100 mmol phosphate buffer, 5 mmol L-cystein, 5 mmol EDTA and 140 μg/ml papain (P4762, Sigma-Aldrich)) at 60 °C. DNA was quantified using the Qubit Quantification Platform (Invitrogen) with the high sensitivity assay, according to the manufacturer’s instructions. Hydroxyproline content as a measure for collagen was quantified using a chloramine-T assay (Huszar et al., 1980) with trans-4-hydroxyproline (H54409, Sigma-Aldrich) as reference. Absorbance values were measured at 550 nm using a plate reader (Synergy^™^ HTX, Biotek) and standard curve absorbance values were used to determine hydroxyproline content in the samples.

#### 2.5.5 Matrix mineralization

Bioreactors were scanned and analyzed with a *μ*CT100 imaging system weekly from day 14 to day 42. Scanning was performed at an isotropic nominal resolution of 17.2 *μ*m, energy level of 45 kVp, intensity of 200 *μ*A, integration time of 300 ms and with twofold frame averaging. To reduce part of the noise, a constrained Gaussian filter was applied with a filter support of 1 and a filter width sigma of 0.8 voxel. Filtered images were segmented to detect mineralization at a global threshold of 24% of the maximum grayscale value. Unconnected objects smaller than 30 voxels were removed through component labeling. Mineralized volume was computed for the total scaffold and the channel region using the scanner manufacturer’s image processing language (IPL) (Bouxsein et al., 2010).

#### 2.5.6 Cell and tissue organization

A quarter of each scaffold (*N* = 4 per condition per time point) was fixed in 3.7% neutral buffered formaldehyde. Samples were washed in PBS, permeabilized for 30 min in 0.5% Triton X-100 in PBS and stained for 1 h with 0.1 *μ*g/ml DAPI, 100 pmol Atto 647-conjugated Phalloidin and 1 *μ*mol/mL CNA35-mCherry (Aper et al., 2014) to visualize cell nuclei, F-actin and collagen, respectively. Samples were subsequently washed and imaged in PBS. Z-stacks of the channel wall were acquired with a confocal laser scanning microscope (Leica TCS SP8X, 20x/0.40 HC PL Fluotar L objective). Z-stacks were deconvolved using the CLME deconvolution algorithm with the Batch Express function of Huygens Professional (version 20.04, Scientific Volume Imaging, The Netherlands). Cell nuclei at the channel wall were subsequently quantified after segmentation using the Huygens Object Analyzer tool (Scientific Volume Imaging) and normalized for the tissue volume. Z-stacks were converted to maximum intensity projections for presentation using FiJi (35). In addition, phalloidin maximum intensity projections were used for cell orientation analyses using a previously described algorithm (Jonge et al., 2013) in MATLAB (version 2019b, The MathWorks Inc., Natrick, MA, USA). In short, the principal direction of each pixel with an actin fiber was derived after eigenvector decomposition of the Hessian matrix. A two-term Gaussian distribution (for bimodal distributions present in the data) was subsequently fitted to the derived actin fiber distribution per sample.

### 2.6 Statistical analyses

Statistical analyses were performed, and graphs were prepared in GraphPad Prism (version 9.3.0, GraphPad, La Jolla, CA, USA) and R (version 4.1.2) (R Core Team, 2020). Data were tested for normality in distributions and equal variances using Shapiro-Wilk tests and Levene’s tests, respectively. When these assumptions were met, mean ± standard deviation are presented, and to test for differences, a two-way ANOVA was performed followed by Turkey’s post hoc tests with adjusted p-value for multiple comparisons. Other data are presented as median ± interquartile range and were tested for a difference between static and dynamic with non-parametric Mann–Whitney U test for each time point with adjusted p-value for multiple comparisons. A *p*-value of <0.05 was considered statistically significant.

## 3. Results

### 3.1 Computational fluid dynamics simulation

From the *μ*CT analyses, a similar pore size distribution for four regions of interest was found with an average pore size of 103 ± 40 *μ*m (Figure S1). Based on the CFD model for the eight RVEs at the micro-scale, the average permeability of the whole scaffold was 4.92×10^-10^ m^2^ (Figure 2B). According to the macro-scale CFD model, the fluid shear stress at the channel wall was uniformly distributed along the longitudinal direction of the channel (*i.e*., from top to bottom) with an average value of 10.62 mPa (maximum WSS = 20.67 mPa, minimum WSS = 9.33 mPa) (Figure 3). Moreover, from the CFD model with idealized cubic pore shape elements the average velocity along the channel was 1.2 mm/s, while the average velocity in the radial direction (*i.e*., across the channel) was 2.3×10^-3^ mm/s (supplementary information, Section 2). Thus, the fluid velocity along the channel was > 520 times higher than the fluid velocity in the radial direction, confirming the assumption that fluid flow through the interface can be neglected. Furthermore, the majority of the fluid flow went through the channel rather than through the scaffold region: the average z-component (*i.e*., in longitudinal direction) of the fluid velocity vector was 1.20 mm/s in the channel compared to 0.16 mm/s in the scaffold region (Figure S2).

**Figure 3.**
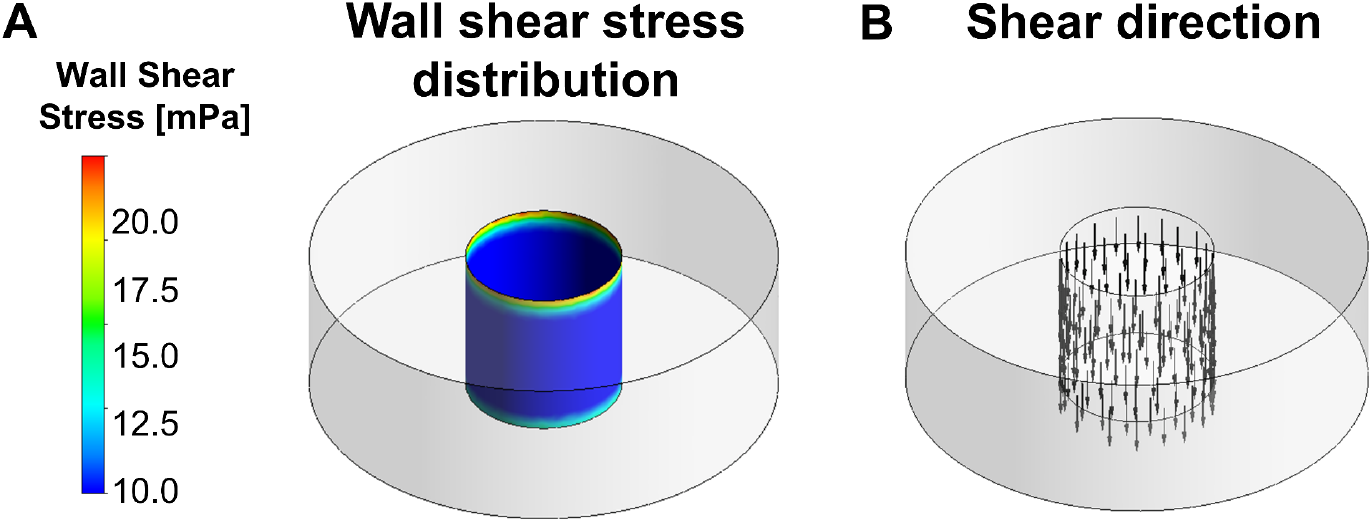
(**A**) Wall shear stress distribution on the inner channel wall of the scaffold. (**B**) The direction of the shear vector was uniform and in the longitudinal direction of the channel.

### 3.2 Cell attachment and distribution

After seeding, cells were found within the pores of the scaffold, in the scaffold and at the channel wall (Figure 4A). SEM revealed that cells at the channel wall bridged the scaffold pores (Figure 4B). The cells collectively covered the channel pores (Figure S3). Then, the influence of fluid flow on cell distribution and potentially on cell proliferation at the channel wall was evaluated. After 42 days, dynamically cultured samples seemed to have formed a thicker cell layer at the channel wall (Figure 4C+D). DNA content quantification of the entire scaffold revealed a time dependent decrease in DNA which was different for statically and dynamically cultured scaffolds (Figure 4E). When comparing the two culture conditions per time point, DNA content at day 28 was higher in statically cultured scaffolds than dynamically cultured samples (Figure 4E). Interestingly, DNA content tended to increase from day 0 to day 14 for both conditions, after which it decreased again to values of day 0 for dynamically cultured scaffolds after 28 days and for statically cultured scaffolds after 42 days. To check whether there was a difference in cell number at the scaffold channel wall between statically and dynamically cultured scaffolds, cell nuclei were counted from Z-stacks of the scaffold channel wall (Figure 4F). At the channel wall, no differences between statically and dynamically cultured scaffolds were found. This suggests that the observed thickening in cell layer is the result of ECM production by cells at the channel wall, rather than cell proliferation.

**Figure 4.**
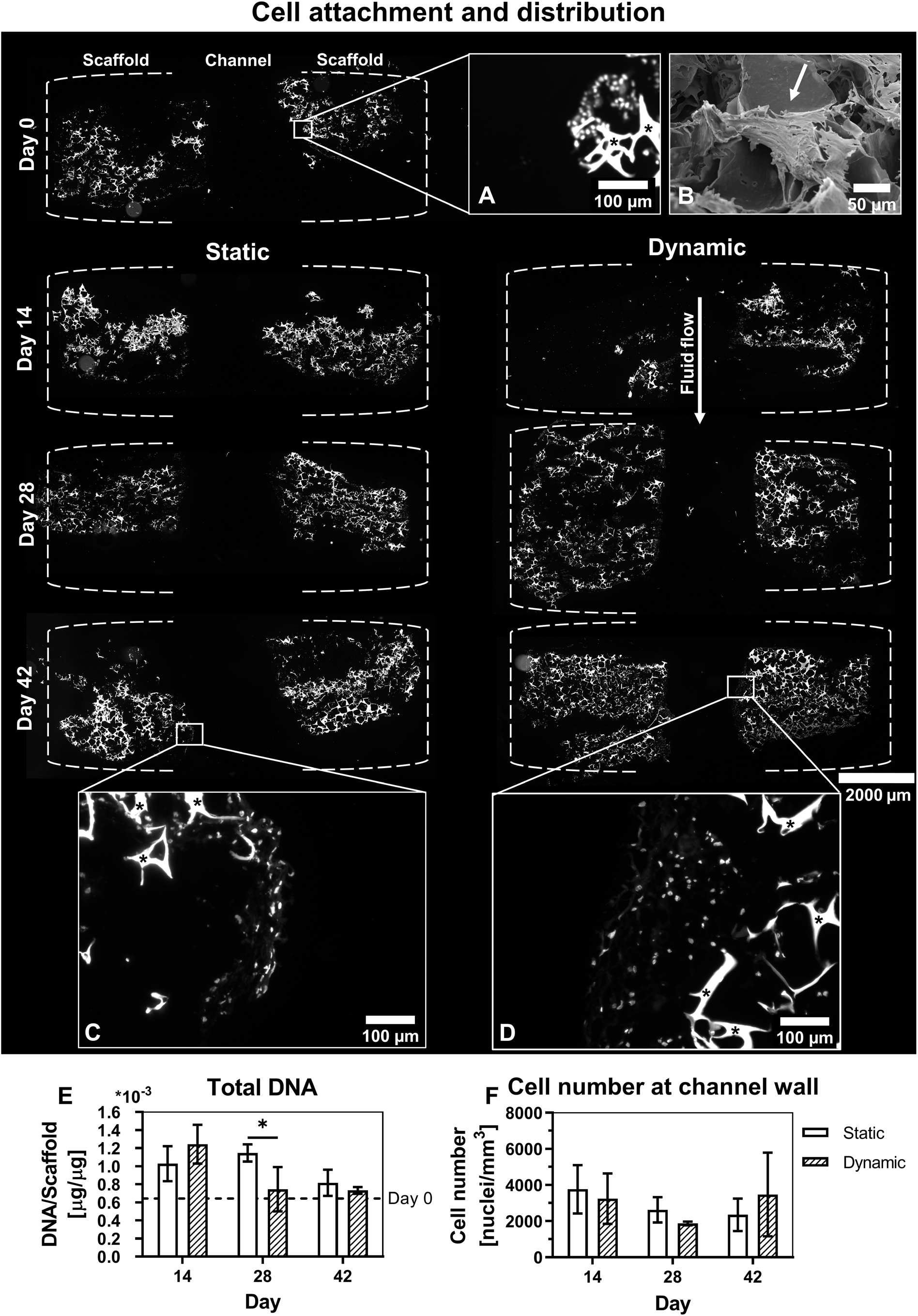
Cell distribution within the scaffold and attachment at the channel wall. (**A**) Scaffold vertical plane sections in which cells and scaffold are visualized with nuclei (DAPI) staining and scaffold autofluorescence (triangular-like structures, indicated with an asterisk) of day 0 samples. (**B**) Cell attachment at day 0 visualized with scanning electron microscopy. (**C**) Cell layer at channel wall for statically cultured scaffolds and (**D**) dynamically cultured scaffolds. (**E**) Total DNA quantification, *p*<0.05 for factor time and time x group interaction (Two-way ANOVA and Turkey’s post hoc tests within each time point), and (**F**) cell count at the channel wall, *ns* (Two-way ANOVA). Asterisks in graphs represent results of post hoc analyses (\**p*<0.05)

### 3.3 Organic matrix growth and mineralization

Collagen deposition was visualized in the horizontal and vertical plane (Figure S4). Vertical plane images revealed collagen formation after picrosirius red staining through the entire scaffold for both statically cultured and dynamically cultured scaffolds (Figure 5A and Figure S4 for more representative images). Collagen content tended to increase with time, with most collagen visible at day 42 of culture for both groups. This was quantified by measuring the hydroxyproline content. Indeed, a time-dependent increase in hydroxyproline content was found (Figure 5C). This increase over time was however similar for statically and dynamically cultured constructs. When zooming in at the channel wall, dynamically cultured scaffolds tended to have a thicker layer of formed tissue than statically cultured scaffolds (Figure 5B). By measuring the thickness of this newly formed tissue in the vertical and horizontal plane, tissue growth at the channel wall could be quantified. Again, a time-dependent increase in tissue thickness was found (Figure 5D). At the channel wall, this time-dependent increase was different for statically and dynamically cultured scaffolds. Dynamically cultured scaffolds had a statistically significant thicker layer of tissue on day 42 at the channel wall than statically cultured scaffolds.

**Figure 5.**
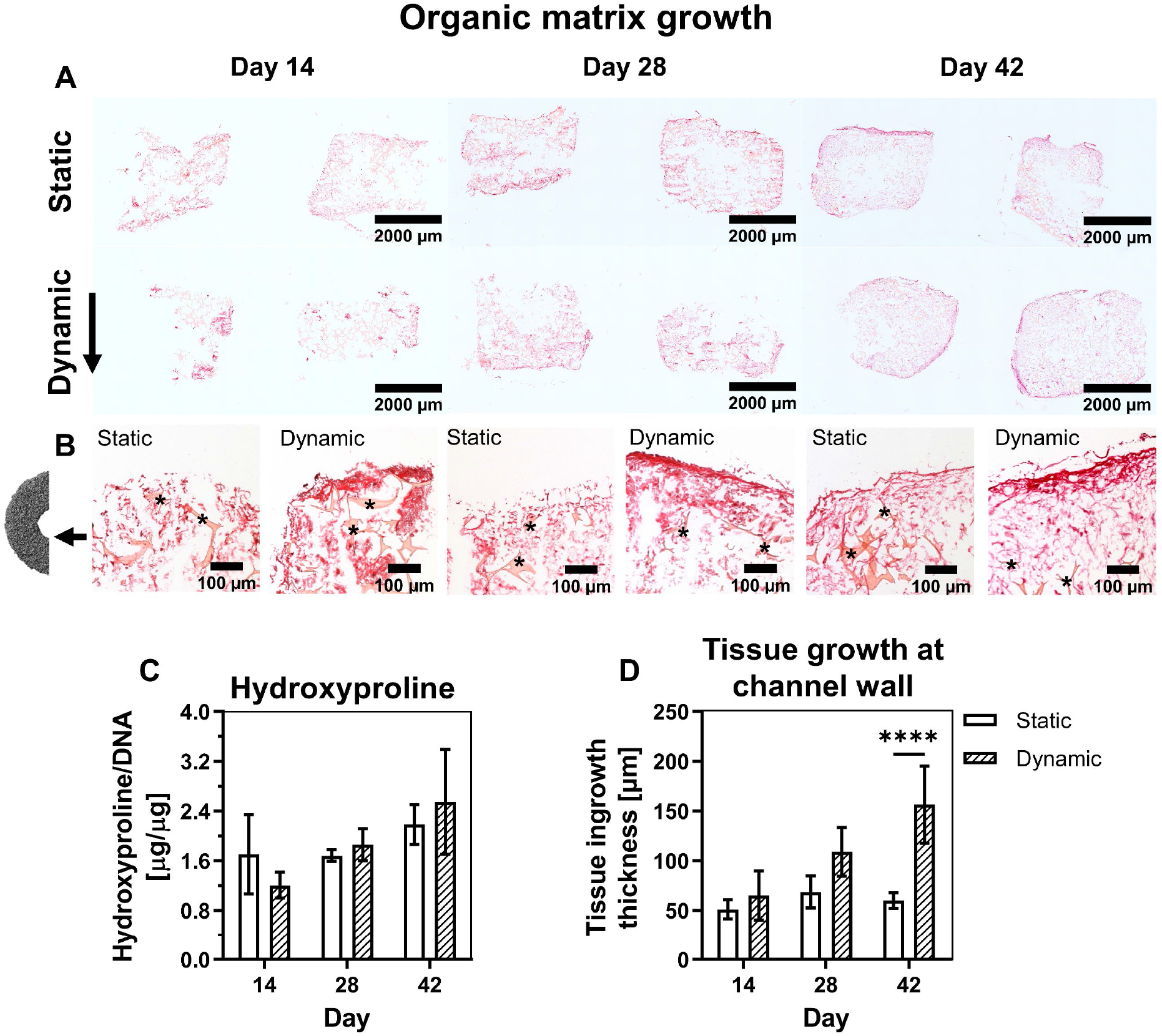
Visualization and quantification of organic matrix growth. (**A**) Micrographs of vertical plane sections stained for collagen (picrosirius red). (**B**) Micrographs of the channel wall of vertical plane sections stained for collagen. Scaffold is indicated with an asterisk (**C**) Hydroxyproline content quantification of the scaffold, *p*<0.05 for factor time (Two-way ANOVA and Turkey’s post hoc tests within each time point) (**D**) Tissue growth quantification at the channel wall, *p*<0.05 for factor time, group, and time x group interaction (Two-way ANOVA and Turkey’s post hoc tests within each time point). Asterisks in graphs represent results of post hoc analyses (\*\*\*\**p*<0.0001).

A positive immunohistochemical staining for collagen type 1 and osteopontin revealed that the formed ECM at the channel wall was of a bone-like character (Figure S5). On day 14, collagen type 1 and osteopontin were mostly present around the cells while on day 42 they were more distributed through the ECM for both statically and dynamically cultured scaffolds. Dynamically cultured scaffolds seemed to have a higher collagen density at the channel wall than statically cultured scaffolds (Figure S5).

Longitudinal *μ*CT monitoring allowed for visualization and quantification of matrix mineralization. From the 3D scans and their quantification, a clear increase in mineralization over time was observed for both groups (Figure 6A+B). Although non-significant, more mineralization seemed present in statically cultured scaffolds. As we were mostly interested in the scaffold channel wall, the scaffold channel volume was also analyzed for the presence of mineralization. Interestingly, at the channel wall differences between statically and dynamically cultured scaffolds could not be observed.

**Figure 6.**
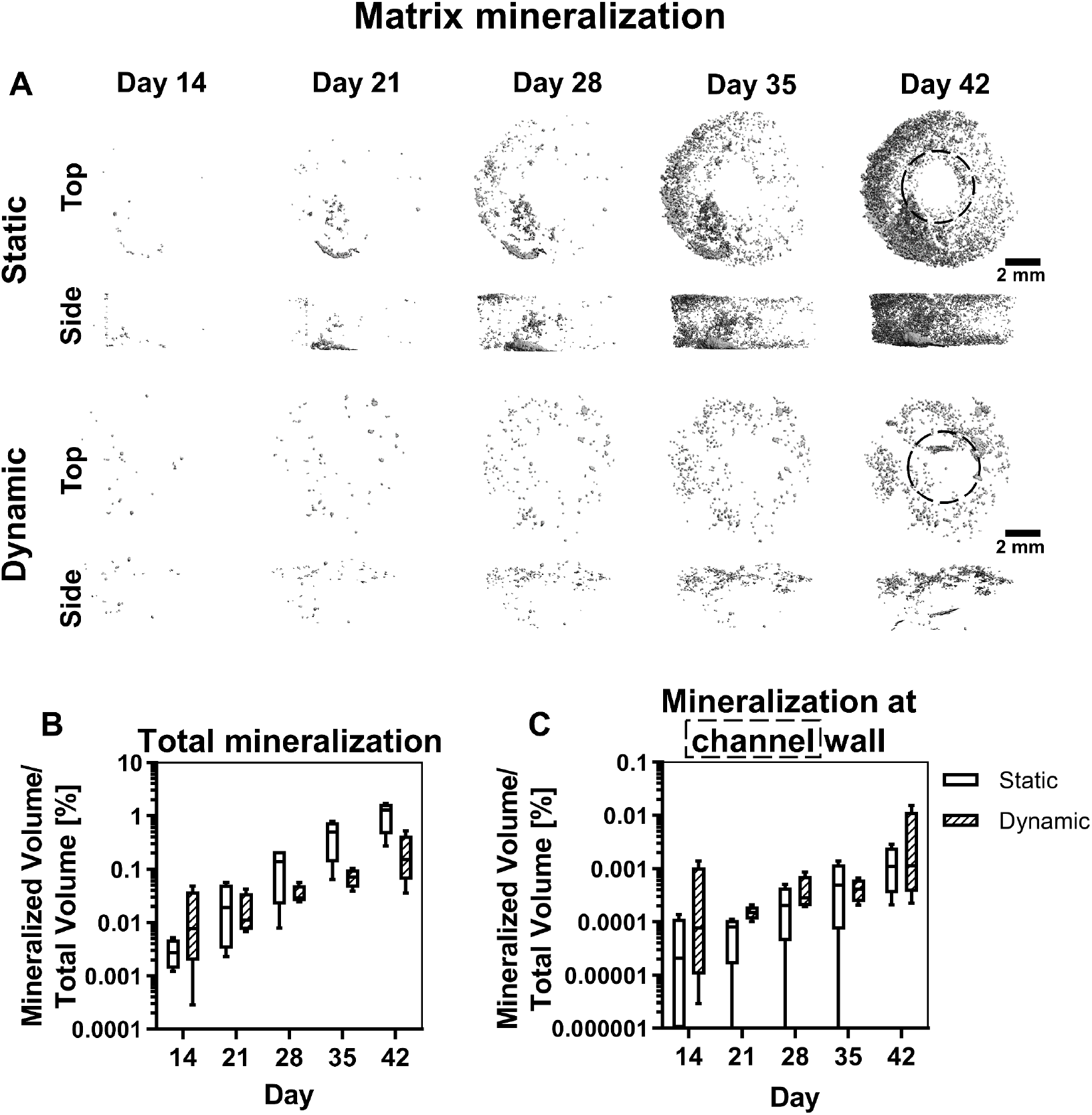
Mineralization over time obtained with *μ*CT-scanning. (**A**) Segmented *μ*CT scans of statically and dynamically cultured scaffolds from day 14 to day 42. (**B**) Mineralized volume of the entire scaffold obtained from *μ*CT scans, *ns* (Mann-Whitney U tests per time point with Bonferroni correction for multiple comparisons). (**C**) Mineralized volume around the channel volume obtained from *μ*CT scans, *ns* (Mann-Whitney U tests per time point with Bonferroni correction for multiple comparisons). Area highlighted with dashed line was analyzed to obtain mineralization at the channel wall. Abbreviations: micro-computed tomography (*μ*CT).

### 3.4 Cell and tissue organization

Over the culture period progression, no clear trend in cell and tissue organization was observed for statically and dynamically cultured scaffolds (Figure 7A+C). From the actin fiber distributions, no consistent influence of directional fluid flow was observed (Figure 7B+D). On day 28, cells tended to align more in the tangential or circumferential direction of the channel for both statically and dynamically cultured scaffolds. This was however not consistent for all scaffolds in the dynamically cultured group.

**Figure 7.**
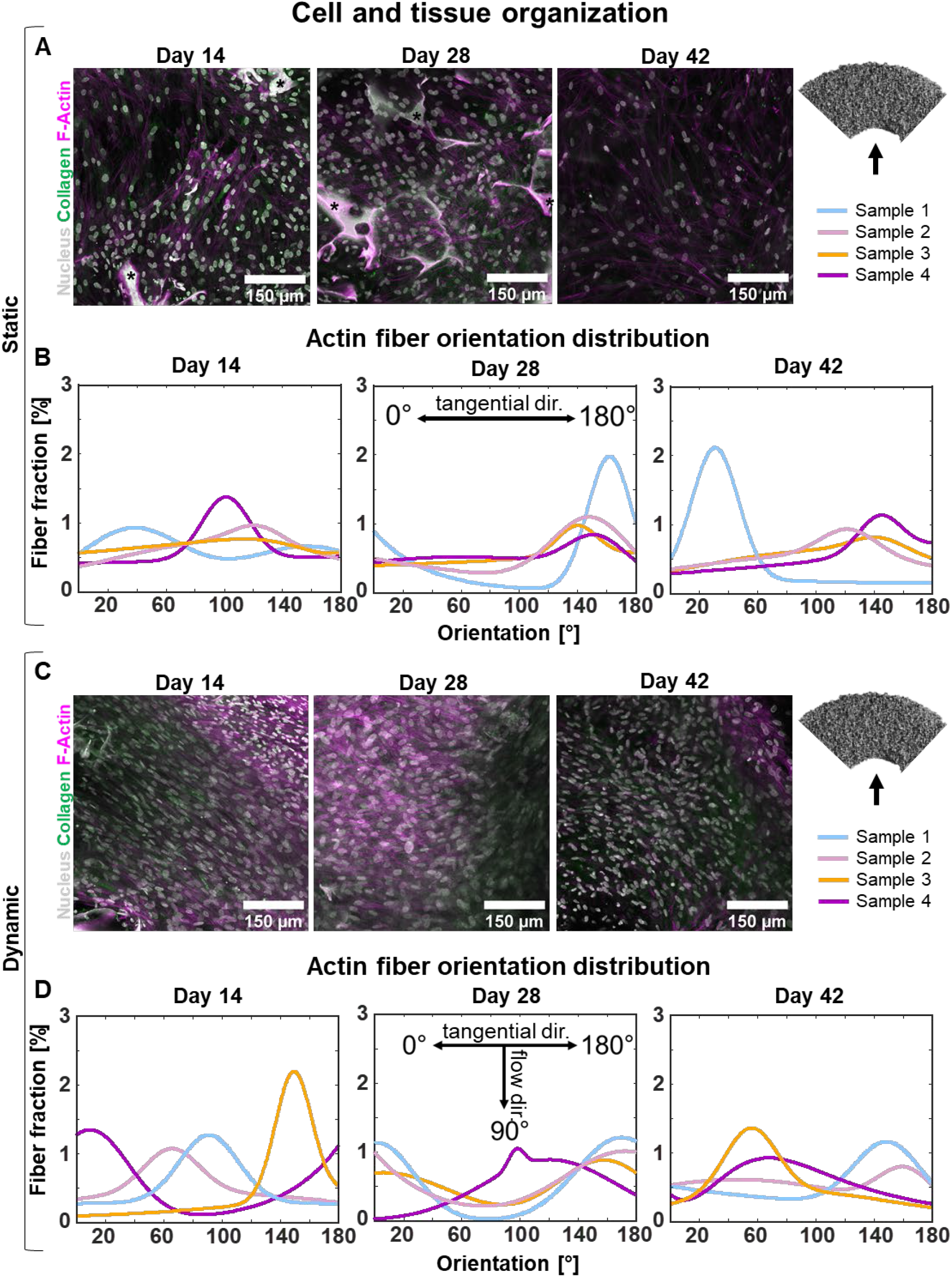
Cell and tissue organization analysis. (**A**) Deconvolved maximum intensity projections of Z-stacks obtained from the channel wall of statically cultured scaffolds. Tissues are stained for the nucleus (gray), collagen (green), and F-Actin (magenta). Scaffold is indicated with an asterisk. (**B**) Gaussian fit (2^nd^ order) of actin fiber orientation distribution of statically cultured scaffolds relative to the tangential direction of the channel and fluid flow. (**C**) Deconvolved maximum intensity projections of Z-stacks obtained from the channel wall of dynamically cultured scaffolds. Tissues are stained for the nucleus (gray), collagen (green), and F-Actin (magenta). (**B**) Gaussian fit of actin fiber orientation distribution of dynamically cultured scaffolds. Abbreviation: direction (dir.).

## 4. Discussion

With the transition in the application of bone tissue engineering strategies from bone regeneration to 3D *in vitro* models, the challenge to create an organized bone ECM has been identified (de Wildt et al., 2019). The creation of an anisotropic and dense bone-like ECM has received little attention and currently available studies were mainly i) performed on 2D substrates over a short period of time, or ii) did not include anisotropy as an outcome.

Although the mechanism by which bone ECM gains its dense and organized structure is not fully understood, curvature (especially concavities), mechanical loading like directional fluid flow, and osteocyte signaling have been identified as potential contributors. In this study, we aimed at evaluating 3D cell and tissue growth and organization in a concave channel with and without directional fluid flow stimulation over a period of 42 days to include the contribution of cell differentiation. As a result, directional fluid flow improved bone-like tissue growth but not organization. After 28 days of culture, when osteogenic differentiation of the cells was likely accomplished, they tended to have a small preference for orienting themselves in the tangential direction of the channel. Even when fluid flow was applied in the perpendicular direction, most samples showed cells with a preference for alignment in the tangential (*i.e*., circumferential) direction of the channel which might be an effect of its curvature (Bidan et al., 2012).

In this study, a CFD model was used to calculate the WSS magnitude and direction. The multiscale model allowed for the calculation of WSS magnitude over the entire channel wall for the highly irregular scaffold used in this study. With an average WSS of 10.62 mPa, mechanical stimulation was within the range of osteogenic and mineralization stimulation for human cells in a 3D environment, based on previous research (Mccoy and Brien, 2010; Melke et al., 2018). To enable this calculation, the assumption that fluid was only flowing in the longitudinal channel direction had to be made. To check whether this assumption was valid, a CFD model was applied on a simplified scaffold geometry comprising of uniform cubical elements as pores. From this model, the made assumption that fluid for generating mechanical stimulation only flows in the longitudinal channel direction seemed valid. In addition, most fluid went through the channel, indicating that within the scaffold culture conditions could be considered static (*i.e*., limited mass transport). This might explain why the difference in collagen formation between statically and dynamically cultured scaffolds were only observed at the channel wall. Another assumption for determining the WSS on the cells was their attachment. Only if cells have a flat attachment to the channel wall, WSS on cells is comparable to the calculated WSS. When cells bridge pores, fluid flow not only induces shear but also strain (Mccoy and Brien, 2010). After seeding, cells indeed bridged the pores at the channel wall. As such, the calculated WSS magnitude might have been an underestimation of the by the cells experienced mechanical load. Cells also covered the channel wall already directly after seeding. It is expected that once they produce ECM, the irregular channel wall gets covered with a more homogeneous tissue layer in which cells experience less strain and are stimulated by mostly WSS (Hadida and Marchat, 2020). However, substantial tissue growth in the channel will likely also change the fluid flow induced shear stress (Zhao et al., 2020). Thus, the fluid flow induced mechanical load is expected to change over time which hinders the interpretation of the obtained results and therefore is a limitation of the present study. In this study, tissue growth and mineralization were already monitored over the entire culture period. Future studies would benefit from including these tissue growth and mineralization parameters in their models to get a more realistic estimation of the change in stress over time and potentially adapt the input flow accordingly (Giorgi et al., 2016; Hadida and Marchat, 2020).

Interestingly, while other studies have reported increased mineralization under the influence of WSS (Akiva et al., 2021; Melke et al., 2018), in our study an opposite effect was observed. Studies with a similar set-up have also found more mineralization in statically cultured scaffolds than dynamically cultured scaffolds (Vetsch et al., 2016; Vetsch et al., 2017). In the used perfusion bioreactor set-up, only half of the medium volume can be replaced whereas in statically cultured bioreactors all the medium can be replaced. To account for this, osteogenic supplements were added in a double concentration under the assumption that they are either consumed or degraded before the next medium change. However, this way of medium replacement might also induce a difference between the groups in protein concentration derived from FBS or in soluble factors produced by the cells (Vis et al., 2020). Recently, the impact of alkaline phosphatase in FBS on mineralization has been shown (Ansari et al., 2022). We therefore suggest that the difference in mineralization is attributed to the bioreactor system and its practical limitations, something that needs to be considered for future experiments using this bioreactor system. At the channel wall, differences in mineralization between statically and dynamically cultured constructs were absent. This suggests that in dynamically cultured constructs, mechanically stimulated cells at the channel wall contributed more to mineralization than cells within the scaffold that likely sensed no to limited shear stress.

In our effort to improve cell and tissue organization in 3D, directional fluid flow was applied in a concave channel. Fluid flow has been shown to stimulate cellular alignment in 2D (Xu et al., 2018), while (mainly concave) curvature has been shown to induce anisotropic collagen formation in 3D (Bidan et al., 2012; Bidan et al., 2013; Callens et al., 2020a). By applying fluid flow in the longitudinal direction of a concave channel in a 3D scaffold, we attempted to identify the main driver of 3D cell and tissue organization. When cells were oriented in the longitudinal direction of the channel, curvature was considered to be neglectable, while if cells aligned in the tangential or circumferential direction, curvature was −0.67 mm^-1^. Over the entire culture period, no clear influences of curvature nor directional fluid flow were observed. Only a small preference for the tangential direction (*i.e*., the channel curvature) was observed after 28 days for both statically and dynamically cultured scaffolds. Fluid flow does not seem to have a consistent influence on cell and tissue organization in the presence of curvature in 3D. Reasons why the influence of curvature was limited in this study could be i) the irregular channel wall, ii) the differentiation state of the cells, and/or iii) the channel diameter and thus the curvature magnitude. First, while in this scaffold, the smallest possible pores were produced to maximize the channel to pore size ratio, scaffold pores might still have induced small and local changes in curvature which could have locally influenced initial cell orientation. This might also explain why after 28 days a small influence of curvature was visible, as once cells have formed a monolayer and produced their own ECM, curvature becomes a more dominant factor than scaffold properties (Kommareddy et al., 2010). However, one would then also expect to see cell alignment in constructs cultured for 42 days which could not be detected in this study. Second, previous research has shown that undifferentiated hBMSCs prefer to avoid curvature and would therefore align in the longitudinal direction of the channel (Callens et al., 2020a). Therefore, cells might have changed their orientation during their differentiation process. Third, bone-like tissue growth is mainly stimulated with higher concavities (Vetsch et al., 2016). However, to avoid closing of the channel, which would have induced unpredictable flow patterns, a trade-off between curvature and channel diameter had to be made.

Recently, prostaglandin E2 signaling by osteocytes was identified as a potential inducer of osteoblast alignment (Matsuzaka et al., 2021). In this study, osteocytes might have been differentiated from hBMSCs but most likely only under influence of mechanical stimulation and towards the end of the culture period (Akiva et al., 2021). The contribution of osteocytes to ECM anisotropy in bone complicates the investigation of organized ECM formation *in vitro,* as in our approach it requires long-term experiments. Osteoblast-osteocyte co-cultures might be performed to overcome this limitation. In addition, the field would benefit from controlled experiments in which multiple cues (*e.g.,* strain, fluid flow, curvature, presence of osteocytes) can be assessed in a high-throughput fashion. Such experiments could lead to the identification of the driving cues for bone ECM growth and anisotropy. Nevertheless, the here presented longitudinal characterization of cell and tissue growth and organization under influence of curvature in the absence and presence of directional fluid flow underlines the complexity of the *in vitro* creation of a dense and anisotropic bone-like ECM, which is desired for *in vitro* bone models (de Wildt et al., 2019).

## 5. Conclusion

In the present study, we presented a computationally informed 3D model for bone-like tissue growth. In our attempt to improve ECM density and anisotropy, cell organization and tissue growth were evaluated under influence of curvature with and without the application of directional fluid flow. Based on the results obtained within this study supported by existing literature, we believe that anisotropy in 3D might be guided by curvature while ECM growth can be improved with the application of WSS. As such, an attempt was made to improve the resemblance of *in vitro* produced bone-like ECM to the physiological bone ECM.

## Author Contributions

BdW, FZ, IL, KI and SH contributed to conception, methodology and design of the study. BdW performed the experiments and analysed the experimental results. FZ and IL performed the computational simulations. BdW and FZ contributed to the figures presented in the manuscript. BdW wrote the original draft of the manuscript. All authors contributed to manuscript revision and approved the submitted version. BdW, FZ, BvR, KI, and SH contributed in the supervision. SH acquired funding for this research.

## Funding

This work is part of the research program TTW with project number TTW 016.Vidi.188.021, which is (partly) financed by the Netherlands Organization for Scientific Research (NWO).

## Conflict of interest statement

The authors declare that the research was conducted in the absence of any commercial or financial relationships that could be construed as a potential conflict of interest.

## Data availability statement

Data can be shared upon reasonable request.

## Ethics statement

Ethical review and approval was not required for the study on human participants in accordance with the local guidelines and institutional requirements. Cell donor provided written informed consent.

## Notes

### Competing Interest Statement

The authors have declared no competing interest.

