## Supplementary Information for "Characterization of three-dimensional bone-like tissue growth and organization under influence of curvature and directional fluid flow"

### 1. Scaffold pore size analyses

To determine the pore size and pore size distribution, the image background was filled with largest possible spheres of which the diameter was derived. From the micro-computed tomography ($\mu$CT) analyses, a comparable pore size distribution for four regions of interest was found with an average pore size of 103 ± 40 $\mu$m (Figure S1). The average pore size was used for the regularized element computational fluid dynamics (CFD) model (Figure S2).

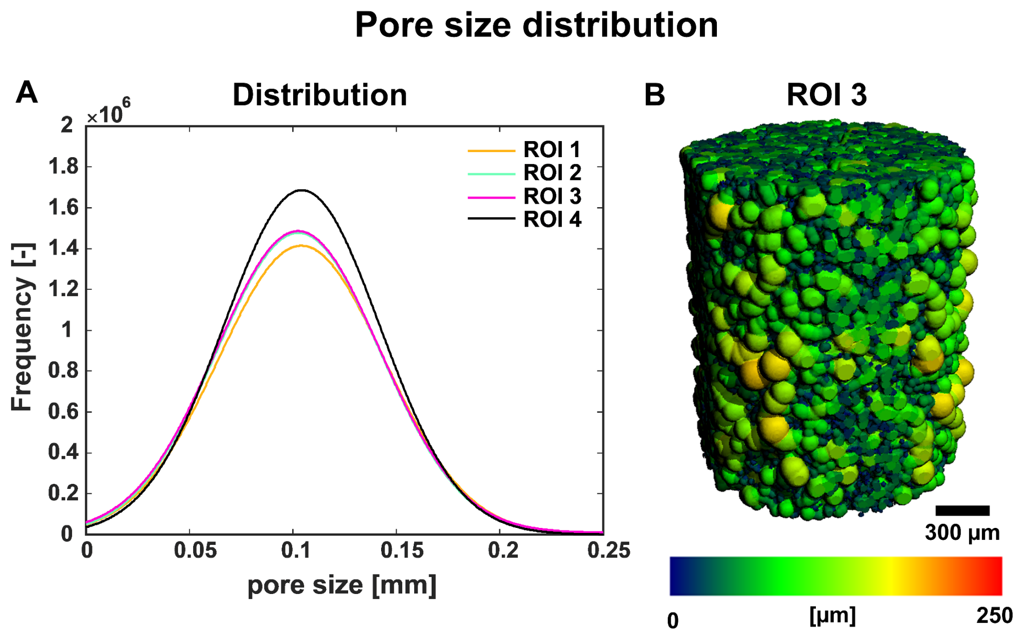


**Figure S1.** (**A**) Gaussian fits of pore size distributions of four different regions of interest (ROI) in the scaffold. (**B**) Pore size distribution was obtained by filling the micro-computed tomography ($\mu$CT) scan background with largest possible sphere per pore.

### 2. Regularized element CFD model

In the macro-scale model, the assumption was made that no fluid flows through the interface of the channel and the scaffold. To check this assumption, a CFD model was evaluated in which a scaffold with idealized cubic pore shape, a porosity of 90% and a pore size of 103 μm was used (Figure S2A). To save the computational costs, representative volumetric elements (RVEs) were modelled with the side faces set as periodic boundaries (Figure S2A). A fluid velocity of 393 μm/s (corresponding to a flow rate of 1.5 ml/min) and a relative pressure of 0 Pa were prescribed at the inlet and outlet (Figure S2A). The other surfaces (*i.e.* struts surfaces) were defined as non-slip walls. The physical properties of flow, mesh strategy and convergence criteria were kept the same as those in the model in Section 2.3.2.

It was found that average velocity along the channel (in Z direction in Figure S2B) was 1.2 mm/s, while the average velocity in the radial direction (*i.e.* across the channel) was $2.3 \times 10^{-3}$ mm/s. Thus the fluid velocity along the channel is > 520 times higher than that the fluid velocity in the radial direction. Moreover, majority of the fluid flow went through the channel rather than through the scaffold region (*i.e.* average Z-direction velocity magnitude = 1.20 mm/s in channel vs 0.16 mm/s in the scaffold region) (Figure S2C).

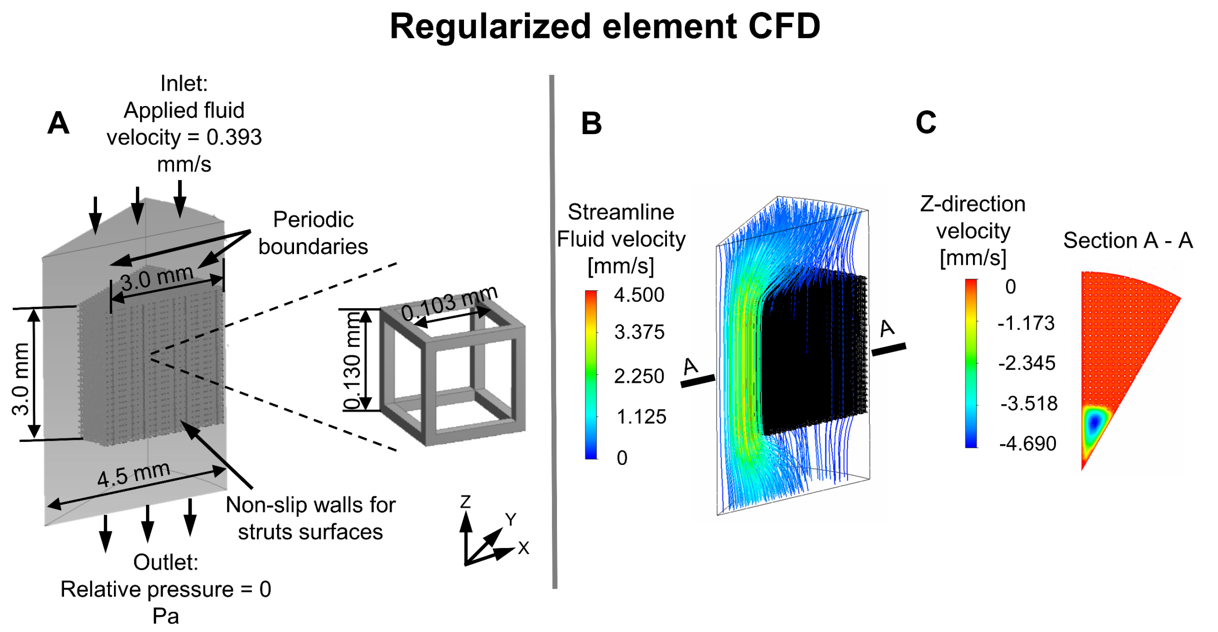


**Figure S2.** (**A**) One representative volumetric element (RVE) of computational fluid dynamics (CFD) model that is based on the scaffold assembled by repeating idealized cubic pore units. (**B**) Overall fluid velocity distribution within the channel and porous scaffold area. (**C**) Z-direction fluid velocity within the channel and porous scaffold (viewed from cross section A-A).

3. **Supporting information on biological experiment**

Cell attachment at the channel wall was assessed at day 0 with scanning electron microscopy. Cells at the channel wall seemed to bridge the scaffold pores and seemed to collectively cover the channel pores (Figure S3).

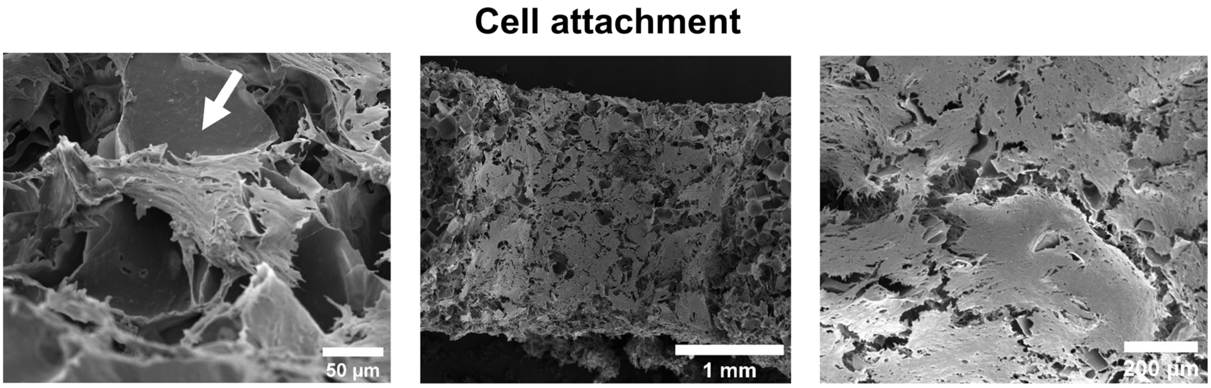


**Figure S3.** Cell attachment at day 0 visualized with scanning electron microscopy. White arrow indicates a cell bridging the scaffold pore.

To visualize collagen deposition, cryosections were prepared in two different planes ($N$ = 4 scaffolds per condition, time point, and plane) and stained with Picrosirius Red. To capture the entire section, tile scans were made with a bright field microscope (Figure S4).

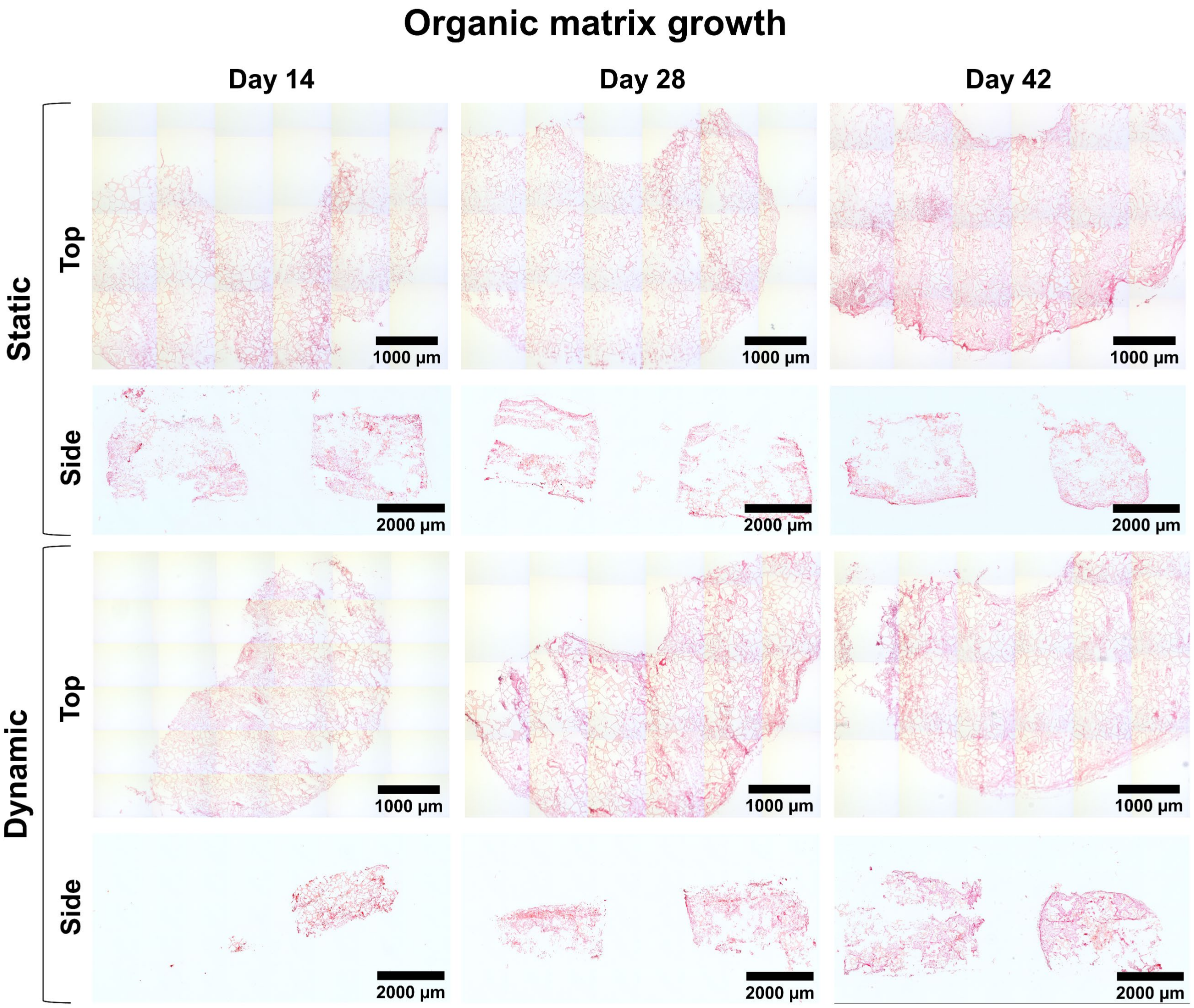


**Figure S4.** Micrographs of vertical plane and horizontal plane scaffold sections stained for collagen (picrosirius red). These images were used for tissue growth quantification.

To check whether the produced extracellular matrix was of a bone-like nature, cryosections (*N* = 4 scaffolds per condition per time point) were prepared and stained with the bone ECM markers osteopontin and collagen type 1. A positive immunohistochemical staining for collagen type 1 and osteopontin revealed that the formed ECM at the channel wall was indeed of a bone-like character (Figure S5). On day 14, collagen type 1 and osteopontin were mostly present around the cells while on day 42 they were more distributed through the ECM for both statically and dynamically cultured scaffolds. Dynamically cultured scaffolds seem to have a higher collagen density at the channel wall than statically cultured scaffolds (Figure S5C+F).

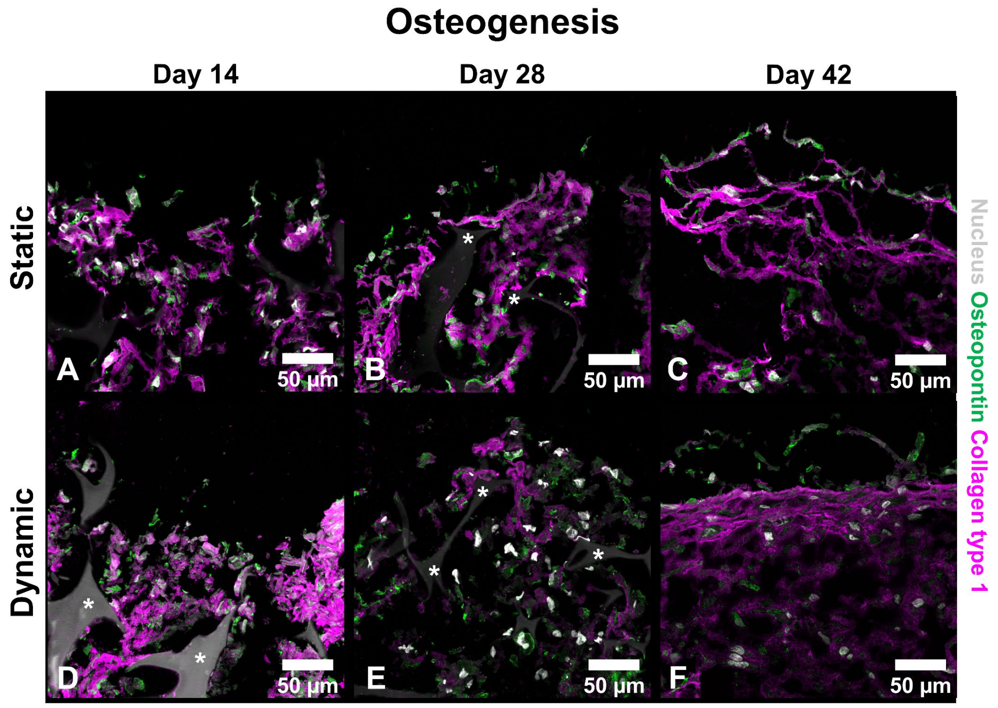


**Figure S5.** Osteogenic differentiation of hBMCSs over time. (**A-F**) Immunohistochemical analysis of sections for collagen type 1 (magenta), the nucleus (gray) and osteopontin (green). Asterisks indicate the scaffold. Abbreviations: human bone marrow-derived mesenchymal stromal cells (hBMSCs).
